# Evaluation of direct grafting strategies in Expansion Microscopy

**DOI:** 10.1101/696039

**Authors:** Gang Wen, Marisa Vanheusden, Aline Acke, Donato Vali, Simon Finn Mayer, Robert K. Neely, Volker Leen, Johan Hofkens

## Abstract

High resolution fluorescence microscopy is a key tool in the elucidation of biological fine-structure, providing insights into the distribution and interactions of biomolecular systems down to the nanometer scale. Expansion microscopy is a recently developed approach to achieving nanoscale resolution in optical imaging. In the experiment, biological samples are embedded in a hydrogel, which is isotropicaly swollen. This physically pulls labels apart, allowing more of them to be resolved. However, in the gelation and swelling process, two factors combine to reduce the signal in the final image; signal dilution and the polymerization reaction, which can damage some fluorophores. Here, we show a chemical linking approach that allows covalent grafting of biomolecular target and reporter in expansion microscopy. Through the combination of a targeting ligand, a reporter moiety and a polymerizable group in a single linker, complex constructs can be prepared in a single, labelling step. We show application of this new series of molecules in the targeting of the cell cytoskeleton, a first example of lipid membranes in expansion microscopy; direct immunostaining with primary and secondary antibodies, and direct grafting of ISH probes and signal amplification initiators (HCR and RollFISH). Our probes allow direct, multiplexed targeting of the cellular blueprint and enable a range of novel imaging approaches in combination with expansion microscopy.

## Introduction

In 2015, Boyden and coworkers introduced expansion microscopy (ExM) as an alternative to super resolution fluorescence microscopy (SRFM) where, instead of engineering the microscope, the specimen is engineered to allow its physical expansion ^1–3^. In the ExM experiment, a fixed sample is permeabilized and structures of interest are labeled with the same fluorescent labeling approaches often used in SRFM such as immunofluorescence or fluorescence in situ hybridization. Next, a chemical crosslinking moiety is introduced, providing protein ^4^ or nucleic acid ^5^ structures with a functional group that is later anchored into a polyelectrolyte polymer meshwork. By infusing the sample with suitable monomers and polymerizing these *in situ*, a swellable hydrogel is formed throughout the sample, which can be expanded upon dialysis in water. Expansion effectively increases the distances between neighboring molecules with a linear expansion factor of ~4.5 fold, yielding an optically-transparent matrix with preservation of the original sample geometry. As such, ExM enables anyone with access to a conventional fluorescent microscope to visualize biomolecules with an effective resolution of ~70 nm.

ExM has been rapidly adopted in the field of fluorescence microscopy, yet it is a relatively young technique and thus under constant evolution. Due to the ability to resolve an increased number of target molecules, ExM has, for example, been combined with multiplexed, error-robust fluorescence in situ hybridization (MERFISH) ^6^ to substantially increase the density of RNAs that can be accurately enumerated in studies of transcription. Furthermore, MERFISH has been demonstrated to be compatible with standard immunofluorescence in ExM, suggesting ExM is an appropriate technique for a multiplexed readout of target molecules of different origin (DNA, RNA, proteins). Due to the intrinsic optical clearing of the sample and thus a higher signal to noise ratio, similar experiments in multicellular structures are readily being performed, shining light on the cell heterogeneity in such a sample. ExM has already lead to the successful imaging of brain tissue slices ^1^’ ^2^’ ^5^’ ^6^, tissue sections of clinical specimens ^7^ and even *Drosophila* tissues.

Despite these exciting results, the transfer to higher resolutions in ExM remains a challenge due to fluorescence signal loss during both the polymerization and digestion steps ^7–11^. Indeed, many fluorophores are prone to degradation during the radical polymerization process, with some being entirely destroyed (e.g. cyanine dyes) ^4^. Furthermore, as sample homogenization (digestion) is crucial to isotropic expansion, fluorescent dyes can be lost due to the random nature and incomplete efficiency of the anchoring step ^12,13^. More specifically, the fraction of fluorophores attached to a proteolytically created protein fragment, that was not also crosslinked to the polymer matrix, are lost during expansion. Finally, upon expansion, the number of fluorescent labels per voxel is diluted by a factor that equals the volumetric expansion (e.g. 4 ^3^ = ~64 fold for the standard ExM protocol ^1^). While this intrinsic dilution of fluorescent labels cannot be prevented, many efforts have been made to bypass the shortcomings of label loss. However, this in turn has led to a multitude of labeling and expansion protocols, all with highly specialized applicability. Hence, experimental parameters and complexity strongly depending on the target molecule and readout scheme chosen. As a result, there is currently no general approach to simultaneously label and image a range of biomolecules (DNA, RNA, protein, membranes).

Here, we demonstrate a direct grafting approach using a tri-functional molecule that enables simultaneous targeting (to a biomolecule of interest), labelling and grafting to the gel. As such, labels are targeted to the relevant location within the cell or tissue and permanently bound at this location throughout the polymerization, expansion and read-out steps. We evaluate the performance of such direct grafting strategies for the singular and multiplexed labeling of proteins, lipid membranes and mRNA which is selectivity achieved through direct grafting of small-molecule ligands, antibodies or *in situ* hybridization probes. Readout can be effected through incorporation of fluorescent dyes, haptens or oligonucleotide reporter barcodes. Pre-polymerization, preexpansion and post-expansion labeling strategies are described and optimized for signal intensity. Furthermore, we show how the different reactivity schemes provide direct access to ISH probes sets as well as signal amplification of low abundance targets through hybridization chain reaction (HCR) ^9^ and RollFISH in ExM. As such, we provide a robust framework for highly multiplexed ExM analysis.

## Results

We have developed a range of ligands, attached to a reporter moiety in a covalent fashion through a dedicated linker. To further ensure covalent attachment of the reporter to the polymeric matrix at the very location of the biological target, a monomer unit (acryloyl) is added to the structure, yielding a trifunctional linker (Figure 1).

**Figure 1.**
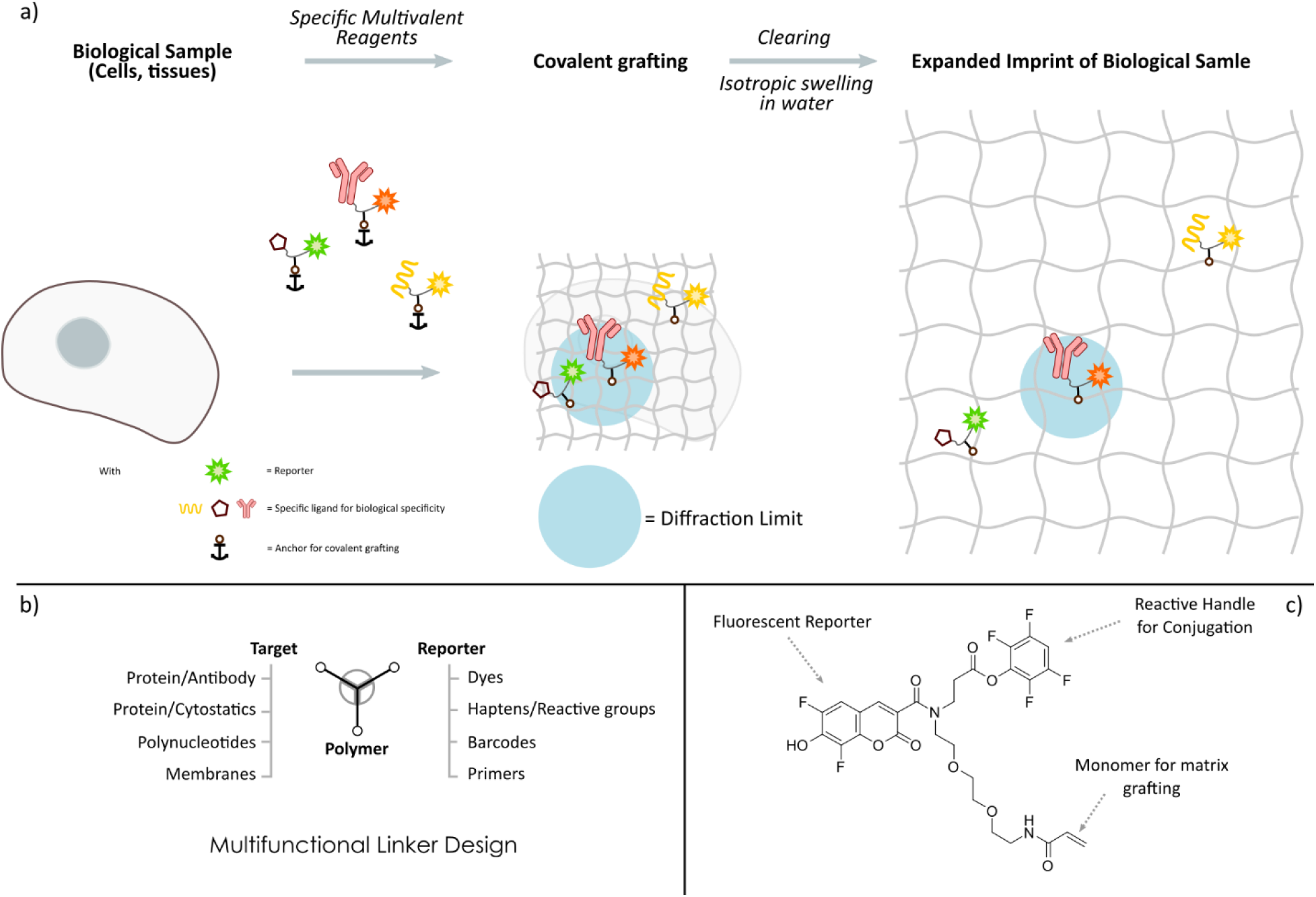
Image depicting the multivalent linker concept and design: a) Multivalent linkers allow for the efficient localization and grafting of signaling moieties within biological samples. b) Through flexible linker design, a range of targets and reports can be obtained. c) Example of direct label trivalent linker with Fluorescent reporter (Pacific Blue), reactive TFP ester for amine conjugation and acrylamide monomer.

In its most simple variant, the reporter moiety is a fluorescent dye. Here, it is imperative to opt for dyes with chromophores which are inert towards the radical polymerization reaction. To evaluate dye stability for use in our multivalent linkers, we screened a large library of commercial and noncommercial organic dyes for stability in the radical polymerization mixture, as measured through fluorescence intensity reduction after polymerization. Here, results are largely in line with earlier observations of Boyden et al. on antibody coupled organic fluorophores (SI) ^4^, where chemical robustness of the chromophore against radical reactions is key to signal survival.

Based on the outcome of these results, a portfolio of the best performing fluorescent dyes in ExM were prepared as multivalent linker, and evaluated for direct fluorescent labeling of cellular structures through coupling to selected ligands for biomolecules (General structure, Figure 1, panel C, full structures in SI).

### Small-molecule targeting of tri-functional labels

As cytoskeletal labeling is both of high biological relevance and often used for evaluating performance in different super resolution approaches, these compounds provided first proving grounds for the approach. Here, labeling density and the size of the label are the two most important parameters to achieve a good super-resolution image. To extend ExM into these studies, we prepared a set of fluorescent expansion linkers, coupling the cytostatic moieties to fluorescent trivalent linkers. For actin staining, a set of phalloidin conjugates were prepared in a single step from the multivalent linkers, with fluorescent reporter dyes spanning the visible spectrum. (Full structures and synthetic procedure in SI). In cellular experiments, these compounds efficiently stain the cytoskeletal actin (Figure 2, panel a-d) in pre-expansion images following standard protocols, and upon expansion. At an expansion factor of ~3.5, this results in an apparent resolution of ~85 nm.

**Figure 2.**
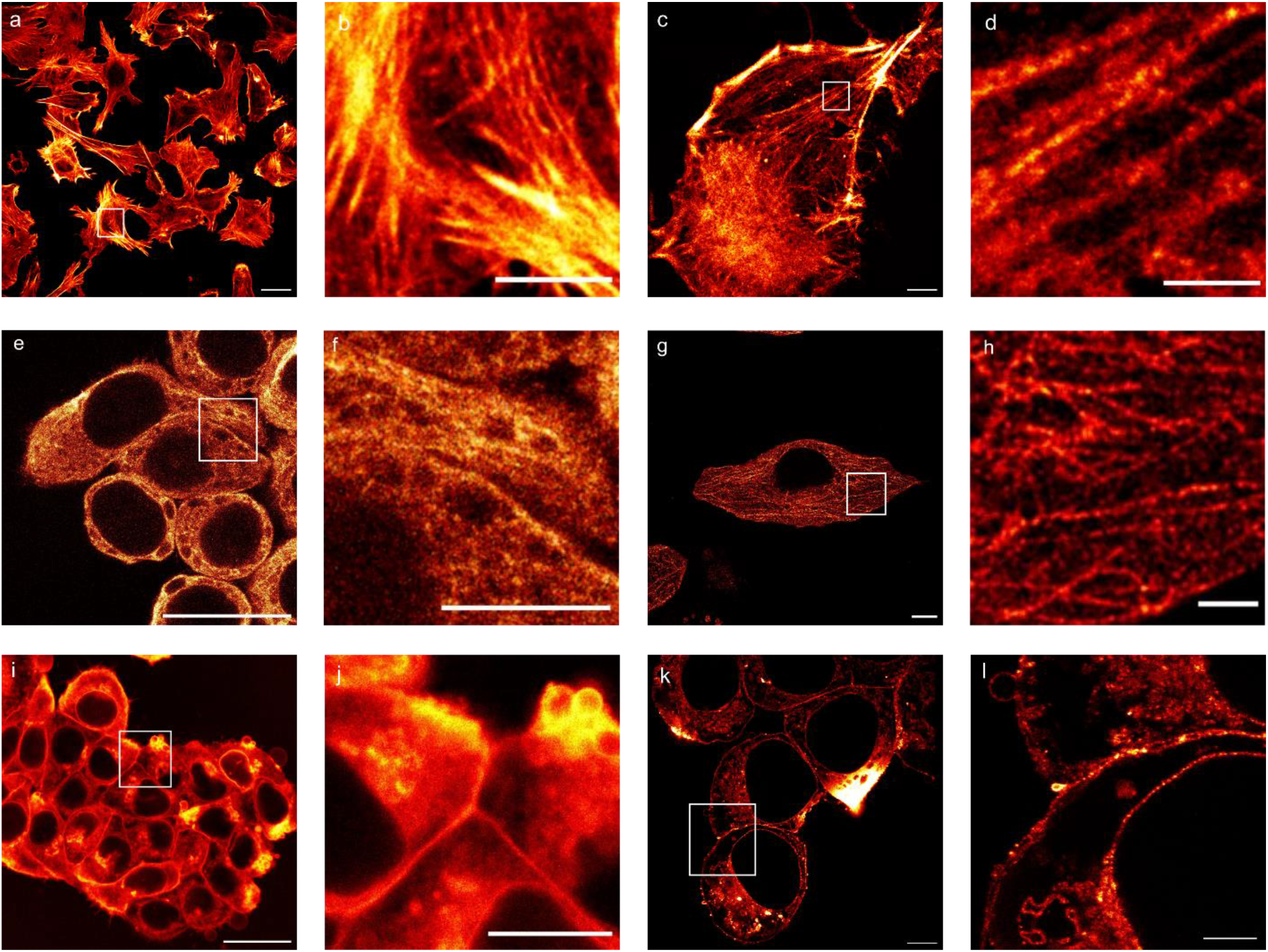
Image showcasing relevant examples of direct grafting in ExM over a range of biological targets: a-d) Direct cytoskeleton staining (Actin) through fluorescent phalloidin conjugated trivalent linker (Rhodamine B, phalloidin, pre- and post-expansion). (a) pre-expansion image with zoom (b) of the boxed area, (c) Average intensity z-projection of post-expansion image with zoom in (d). e-h) Direct immunostaining of alpha-tubulin in ExM though secondary antibody conjugation to fluorescent trivalent linker (Rhodamine 6G, GAM, pre- and post-expansion). (e) pre-expansion image with zoom (f) of the boxed area, (g) post-expansion image with zoom in (h). i-l) Expansion of cellular phospholipid membranes through lipid conjugation to fluorescent trivalent linker (Pacific Blue, DSPE, pre- and post-expansion). i) pre-expansion image with zoom (j) of the boxed area, (k) post-expansion image with zoom in (l). Scale bars: 25 μm (a, c, e, g, i, k), 10 μm (b, d, f, h, j, l).

Furthermore, in what is a first example of lipid membrane expansion microscopy, trivalent fluorescent lipids, carrying DSPE (1,2-Distearoyl-sn-glycero-3-phosphoethanolamine), stain phospholipid bilayers, with the membrane structure and microstructure clearly visible after the expansion process. (Figure 2, panel i-l, structure and further images in SI). This experiment is also a testimony to the overarching concept of substituting a biological structure with permanently-tethered labels, as lipid membranes structures will not survive the expansion process, but their signal is permanently imprinted using our approach.

### Immunostaining using tri-functional labels

Direct coupling of the trivalent systems to antibodies allows single-step immunostaining in expansion microscopy, with both primary and secondary antibodies. To demonstrate this, we imaged HeLa cells which were immunofluorescently labelled for alpha-tubulin via secondary antibodies, coupled to a fluorescent trivalent linker, and this with examples over a broad wavelength range (Structure and synthetic procedures in SI). The typical signal of these tubulin structures is well preserved in both pre- and post-expansion processes (Figure 2, panel e-h, pre- and post-expansion). Furthermore, the use of primary antibodies with a trivalent linker was explored, with conjugates again obtained from a single step reaction. Here, HeLa cells were immunostained for nuclear lamin with a respective anti-lamin A/C primary antibody and this coupled to Pacific Blue. To confirm successful grafting of the primary antibody after coupling to the linker, a secondary antibody staining (GAM-Atto 488) was performed on the same samples, where clear signal overlap in both channels demonstrates retained specificity of the lamin antibody conjugate (SI).

In general, the fluorescent multivalent tags provide excellent staining of their biological targets, with signals well retained after swelling, and this at most common excitation wavelengths. This establishes a general method for providing permanent signatures of fine biomolecular structures, with the possibility to directly generate constructs for a wide range of targets.

As discussed above, the reaction conditions of the polymerization process lead to moderate to near-complete destruction of many of the popular fluorescent dyes. To circumvent this issue, bio-orthogonal reactive groups and/or haptens can be grafted onto the polymer matrix, as “dark” labels prior to polymerization, and subsequently stained with their fluorescent counterparts pre- and post-expansion (SI). The potential of such a post-polymerization labeling step and its impact on brightness in ExM is exemplified by a recent manuscript of Shi and coworkers ^14^.

Although multivalent linkers enable simple and versatile labeling strategies in expansion microscopy, their multiplexing capacity remains limited. By introducing molecular (DNA) barcodes as permanent labels in the polymer network, the method becomes amenable to multiplexed and repeated readout schemes. As such, oligo DNA strands (ODs) hold the potential to serve as hybridization reporter and barcode, and they are readily combined with the multifunctional linkers. After cellular target labeling, the oligo-DNA strands are covalently anchored on a polymeric matrix, leaving a permanent signature of the biological information once present at the location, open to further use for signal reconstruction.

In practice, DNA oligos are attached to the trivalent linker through active ester based coupling with amine terminated oligonucleotides or thiol-maleimide chemistry. These stable conjugates can be directly coupled to e.g. antibodies in single step reactions. In line with previous examples on the staining of a range of biomolecules in expanded gels, we were able to use OD-labeled antibodies (Figure 3) for specific recognition of their respective targets, followed by gelation, expansion and hybridization with fluorescently labeled reporter probes. For example, we effected immunostaining of alpha-tubulin via direct grafting of DNA-conjugated secondary antibody and fluorescent oligo-based readout post-expansion. This is also an example of how post-gelation addition of dyes allows use of e.g. cyanine reporter dyes in ExM, a dye otherwise destroyed in the polymerization process. It should also be noted that this method allows for highly multiplexed imaging, without being inherently restricted to the spectrally resolvable dyes. In addition, this approach allows an assessment of the impact of the labeling scheme, and while rhodamine B direct staining is quite resistant to radical polymerization, post-expansion labeling through a fluorescent readout oligo clearly performs better, likely due to signal loss (of the rhodamine B) during polymerization (approx. 50-60% dye survival). These examples demonstrate that the use of oligo-reporters in combination with the use of primary antibodies has significant potential for multiplexed imaging, where high signal-to-noise is needed.

**Figure 3.**
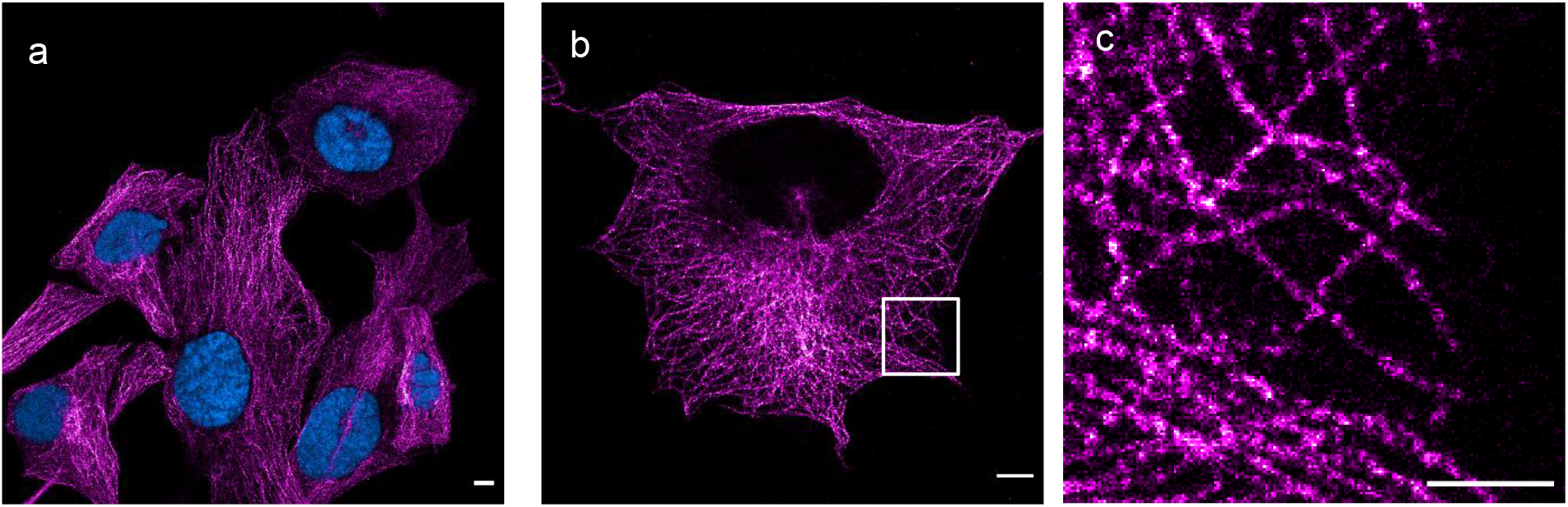
Immunostaining of alpha-tubulin via direct grafting of DNA-conjugated secondary antibody and fluorescent oligo-based readout post-expansion. (a) Two-color image with a nuclear DAPI staining in cyan and immunostaining of alpha-tubulin, visualized with readout oligo 1 (Cy5, sequence in SI) in magenta. (b) same staining as (a), with the DAPI-channel removed to show the high signal to noise ratio and low background staining in the nucleus. (c) Zoom of the boxed area in (b). Scale bars: 10 μm (a-b), 5 μm (c).

The targeting moiety can also be a selective oligonucleotide (ODN). Through the use of fluorescent multivalent linkers, FISH probes for use in ExM can be prepared in a single step from pools of commercial ODN tiling probes. When ODN reporters are combined with ODN targeting moieties, the covalent and stable nature of the ODN reporter allows it to serve as a signal aggregator. In turn, this allows the use of signal amplification approaches in expansion microscopy, and the study of low-abundance species with high signal-to-noise ratios relative to standard approaches. Again, sequential or combined/orthogonal coupling of the multivalent linker to oligo tiling probes with hybridization chain reaction initiator sequence (HCR) ^15–18^or amplification primers (RollFISH) ^19–21^ allows the creation of multifunctional ISH probe sets, in only a few hours and from commercial oligonucleotides.

To demonstrate this, we conjugated tiling probe libraries to the smHCR initiator probes, all coupled through multifunctional linkers (Figure 4, panel a). We evaluated several reactivity schemes, but most convenient examples use orthogonal linker chemistry to ensure correct coupling. Here, coupling chemistry can be effected through amine-to-amine coupling (using TCO-tetrazine cycloaddition as orthogonal to reactive ester-amine coupling). Probe design and targets were in line with literature examples (Shah et al., Wu et al.) for labeling of the gusb transcript in HeLa cells (Experimental information in SI). The images show bright and discrete spots, which are in good agreement with literature examples (Figure 4, panel f-i). In a second approach, RollFISH was implemented in ExM though the coupling of initiator strands to the multifunctional linker. By simply switching out the HCR initiator strand for the RollFISH primer, and in line with literature probe designs, probe libraries were efficiently prepared (Experimental information in SI). Upon addition of reporter oligo, bright and discrete spots were formed (Figure 4, panel b-e), Here, expansion with water was not possible, since the hybridization based signal would not be retained due to DNA de-hybridization, due to charge repulsion of the two strands. This problem has previously been addressed through a secondary polymerization to stabilize the gel prior to higher ionic strength buffer addition ^5^. In our experiments, we tried different buffers to strike a balance between expansion factor and probe retention, and conducted hybridization/expansion experiments in 1X DPBS, 0.5X DPBS, 500 μM MgCl_2_ and 50 μM MgCl_2_. The best results were obtained with 500 μM MgCl_2_, still yielding an expansion factor of 4, only marginally different from the typical expansion factor of 4.5 reached with water. 1X DPBS 0.5X DPBS, and 50 MgCl_2_ gave expansion factors of 1.9, 2.4 and 4.5, respectively. As such, and to the best of our knowledge, we provide first examples of signal amplification in ExM where the actual initiator of the amplification process is directly imprinted at the prior location of the biological information, enabling high S/N and repeated read-out schemes. Similarly, minor modifications of the reaction schemes combined with the presented use of direct grafting with multifunctional linkers, should readily lead to the incorporation of several of the other multiplexing/signal amplification methods recently developed (e.g. immuno-SABER ^22^, MERFISH ^6^, …) into the ExM toolbox.

**Figure 4:**
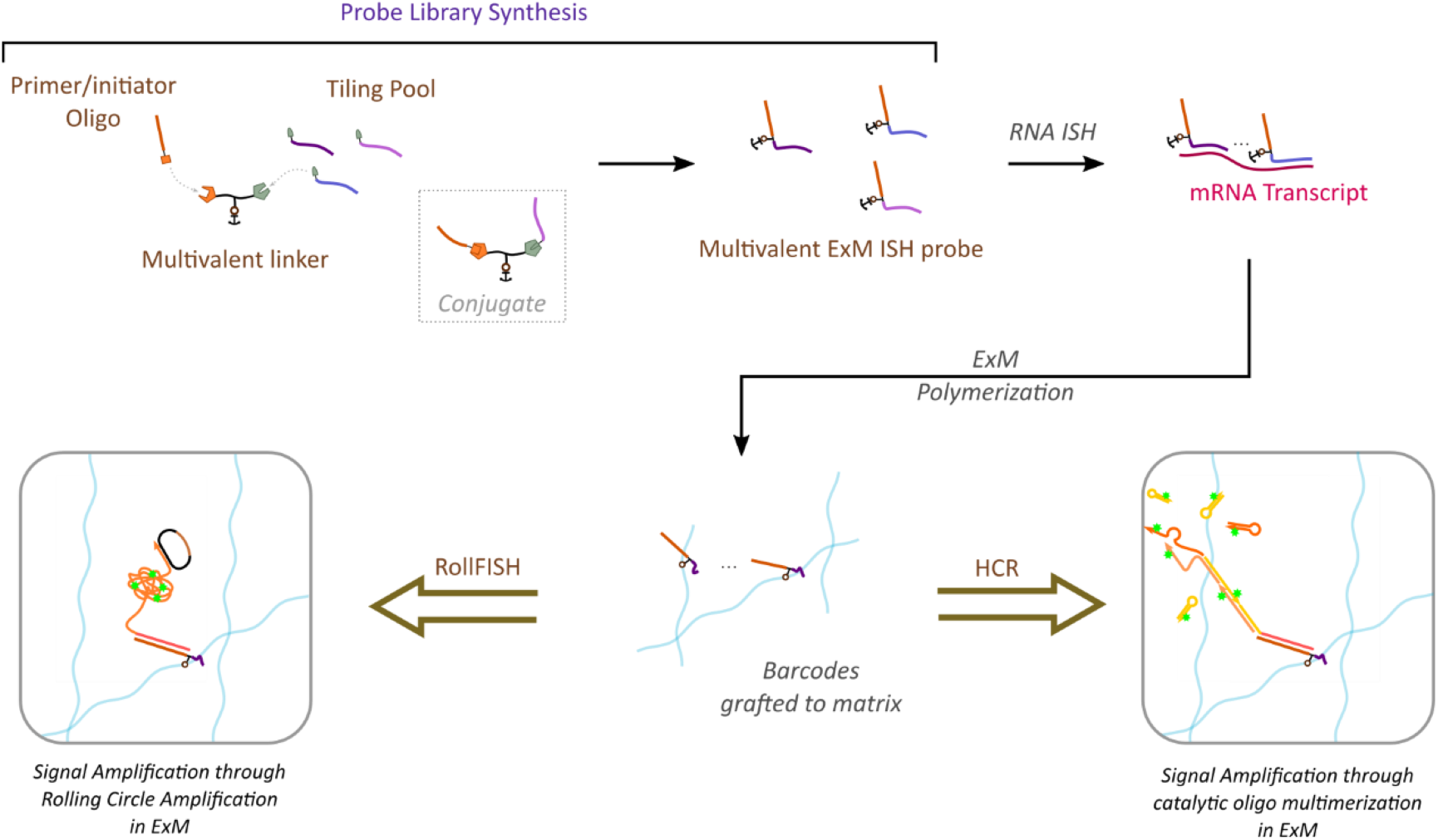
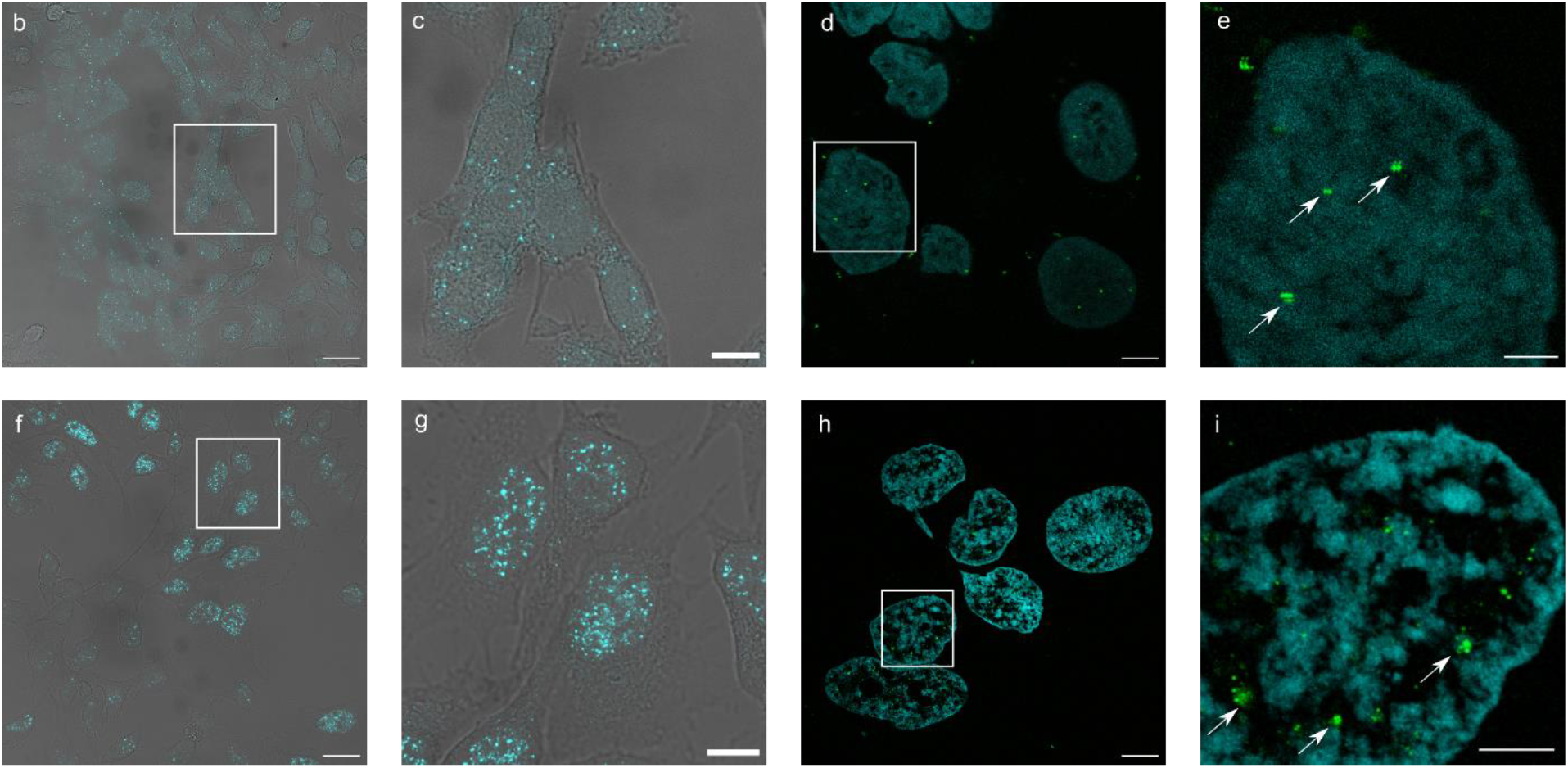
Direct reporter grafting approaches with signal amplification. a). Concept and library synthesis. Convergent synthesis of HCR and ROLLFISH libraries from commercial oligonucleotides and multivalent linker with orthogonal chemistry provides a rapid entry to the construction of ISH probes for use in ExM. b-e) **ROLLFISH in ExM** on regular HeLa cells, (gusb transcript, Alexa Fluor 546 read-out probe, pre and post expansion). (b) pre-expansion image of gusb transcript (cyan) with a bright field overlay (c) zoom of the boxed area in (b), (d) post-expansion image of the gusb transcript (green spots) inside a DAPI stained nucleus (cyan), (e) zoom in of the boxed area in (d) showing defined bright RNA signal spots, as indicated by white arrows. f-i) **smHCR in ExM** on regular HeLa cells trough covalent grafting of initiator strands (gusb transcript, Alexa Fluor 546 read-out probe, pre and post expansion). (f) pre-expansion image of gusb transcript (cyan) with a bright field overlay (g) zoom of the boxed area in (f), (h) post-expansion image of the gusb transcript (green spots) inside a DAPI stained nucleus (cyan), (i) zoom in of the boxed area in (h) showing defined bright RNA spots indicated by white arrows. Scale bars: 25 μm (b, d, f, h), 10 μm (c, e, g, i).

## Conclusion

We describe the preparation of multifunctional small-molecule linkers that allow the direct grafting of biomolecule position and identity to a polymeric matrix, with both parameters preserved after sample clearing and expansion. We have shown how these compounds can be adapted to provide ExM imaging of a range of biological targets, with signal reported through direct fluorescent dyes, fluorescent antibodies and oligonucleotide hybridization. Given the flexibility of the multivalent linkers, and as an unlimited number of oligo reporters can be grafted onto the polymeric matrix in the first step of the ExM protocol, these methods allow simple, repeated and multiplexed readout in ExM, with combinations of the abovementioned methods. In light of the inherent specificity associated with the DNA barcodes and their application in a range of complex biological studies, we foresee rapid extension of direct grafting strategies into complex and highly multiplexed read-out schemes.

## Supporting information

Visualization of DNA-conjugated antibodies using an SDS-PAGE gel assay.

## References

1. Chen, F., Tillberg, P. W. & Boyden, E. S. Expansion microscopy. Science 347, 543–548 (2015).

2. Chozinski, T. J. et al. Expansion microscopy with conventional antibodies and fluorescent proteins. Nat. Methods 13, 485–488 (2016).

3. Ku, T. et al. Multiplexed and scalable super-resolution imaging of three-dimensional protein localization in size-adjustable tissues. Nat. Biotechnol. 34, 973–981 (2016).

4. Tillberg, P. W. et al. Protein-retention expansion microscopy of cells and tissues labeled using standard fluorescent proteins and antibodies. Nat. Biotechnol. 34, 987–992 (2016).

5. Chen, F. et al. Nanoscale imaging of RNA with expansion microscopy. Nat. Methods 13, 679–684 (2016).

6. Wang, G., Moffitt, J. R. & Zhuang, X. Multiplexed imaging of high-density libraries of RNAs with MERFISH and expansion microscopy. Sci. Rep. 8, 4847 (2018).

7. Gambarotto, D. et al. Imaging cellular ultrastructures using expansion microscopy (U-ExM). Nat. Methods 16, 71 (2019).

8. Gao, M. et al. Expansion Stimulated Emission Depletion Microscopy (ExSTED). ACS Nano 12, 4178–4185 (2018).

9. Halpern, A. R., Alas, G. C. M., Chozinski, T. J., Paredez, A. R. & Vaughan, J. C. Hybrid Structured Illumination Expansion Microscopy Reveals Microbial Cytoskeleton Organization. ACS Nano 11, 12677–12686 (2017).

10. Li, R., Chen, X., Lin, Z., Wang, Y. & Sun, Y. Expansion enhanced nanoscopy. Nanoscale 10, 17552–17556 (2018).

11. Wang, Y. et al. Combined expansion microscopy with structured illumination microscopy for analyzing protein complexes. Nat. Protoc. 13, 1869 (2018).

12. Park, Y.-G. et al. Protection of tissue physicochemical properties using polyfunctional crosslinkers. Nat. Biotechnol. 37, 73–83 (2019).

13. Truckenbrodt, S., Sommer, C., Rizzoli, S. O. & Danzl, J. G. A practical guide to optimization in X10 expansion microscopy. Nat. Protoc. 14, 832 (2019).

14. Shi, X. et al. Label-retention expansion microscopy. bioRxiv 687954 (2019). doi:10.1101/687954

15. Dirks, R. M. & Pierce, N. A. Triggered amplification by hybridization chain reaction. Proc. Natl. Acad. Sci. 101, 15275–15278 (2004).

16. Choi, H. M. T., Beck, V. A. & Pierce, N. A. Next-Generation in Situ Hybridization Chain Reaction: Higher Gain, Lower Cost, Greater Durability. ACS Nano 8, 4284–4294 (2014).

17. Shah, S. et al. Single-molecule RNA detection at depth by hybridization chain reaction and tissue hydrogel embedding and clearing. Development 143, 2862–2867 (2016).

18. Choi, H. M. T. et al. Third-generation in situ hybridization chain reaction: multiplexed, quantitative, sensitive, versatile, robust. Development 145, dev165753 (2018).

19. Nilsson, M. et al. Making ends meet in genetic analysis using padlock probes. Hum. Mutat. 19, 410–415 (2002).

20. Wu, C. et al. RollFISH achieves robust quantification of single-molecule RNA biomarkers in paraffin-embedded tumor tissue samples. Commun. Biol. 1, 209 (2018).

21. Strell, C. et al. Placing RNA in context and space – methods for spatially resolved transcriptomics. FEBS J. 286, 1468–1481 (2019).

22. Highly multiplexed in situ protein imaging with signal amplification by Immuno-SABER | bioRxiv. Available at: https://www.biorxiv.org/content/10.1101/507566v1.supplementary-material. (Accessed: 14th June 2019)

