## Supplementary material for "Evaluation of direct grafting strategies in Expansion Microscopy": Visualization of DNA-conjugated antibodies using an SDS-PAGE gel assay.

### General synthetic procedures

Unless otherwise indicated, all solvents and organic reagents were obtained from commercially available sources and were used without further purification. Dry solvents for reaction were used as received from commercial sources. All reaction progress was monitored using thin layer chromatography (TLC) with silica gel plates (Kieselgel 60 F254 plates, Merck) under UV light and LC-MS (Waters Acquity UPLC/ SQD). Mass spectra was obtained using a Waters Acquity UPLC-SQD mass spectrometer. High resolution mass spectra (HRMS) were acquired with a quadrupole orthogonal acceleration time-of-flight mass spectrometer (Synapt G2 HDMS, Waters, Milford, MA). $^{1}$H NMR spectra was recorded on a Bruker Avance 300 MHz or a Bruker Avance Ⅱ+ 600 MHz instrument, and $^{13}$C NMR spectra was recorded at a Bruker 600 Avance Ⅱ+ instrument using DMSO-$d_6$, MeOH-$d_4$ or $CDCl_3$ as a solvent and tetramethylsilane (TMS) as an internal standard.

### General procedure 1: Preparation of multivalent linker and fluorescent dye coupling

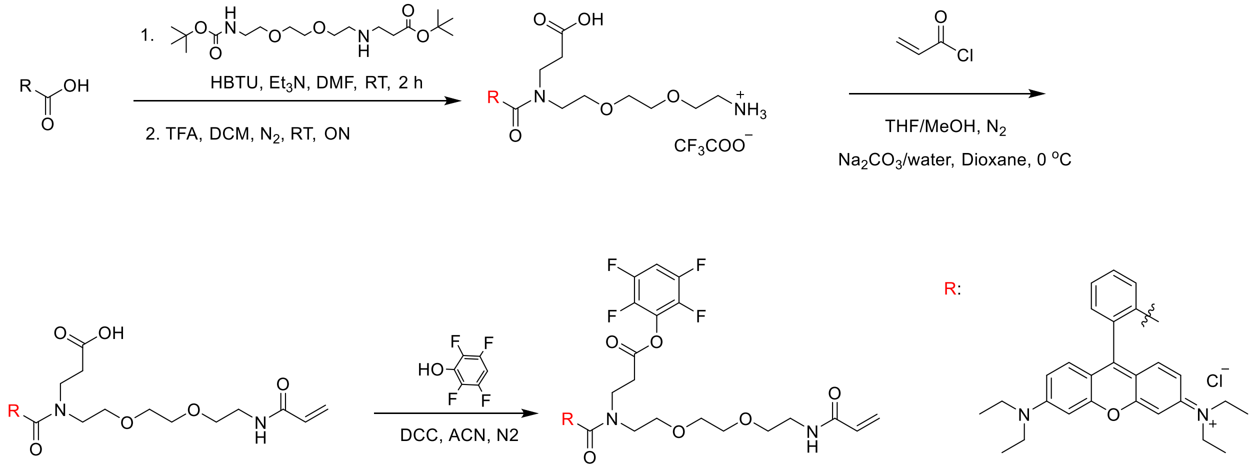


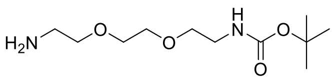

To a solution of 1,2-bis(2-aminoethoxy)ethane (13.1 mL, 89.7 mmol) in dry DCM (80 mL) was added a solution of di-tert-butyl dicarbonate (2.91 g, 15.0 mmol) in dry DCM (25 mL) dropwise at 0 °C over 30 min. The reaction mixture was allowed to warm to room temperature and stirred overnight at the same temperature. After the reaction finished, the solvent was removed under reduced pressure. The residue was dissolved in water (30 mL) and then extracted with DCM (35 mL, 4x). The combined organic phase was dried over $MgSO_4$ and evaporated to get the intermediate **S1** (light yellow oil, 86%). $^{1}$H NMR (300 MHz, $CDCl_3$) δ 5.10 (s, 1H), 3.62 (s, 4H), 3.58 – 3.46 (m, 4H), 3.32 (dd, $J$ = 10.3, 5.2 Hz, 2H), 2.88 (t, $J$ = 5.2 Hz, 2H), 1.45 (s, 9H).

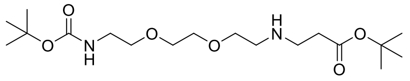

To a solution of **S1**(4.3 g, 17.4 mmol) in EtOH (50 mL) was added tert-butyl acrylate (2.25 g, 17.6 mmol) and triethylamine (2.4 mL, 17.4 mmol), the reaction mixture was stirred overnight at room temperature. After complete reaction, the solvent was removed under reduced pressure and the residue was purified by column chromatography to obtain the desired product **S2** as a light yellow oil (58%) $^1$H NMR (300 MHz, $CDCl_3$) δ 5.19 (s, 1H), 3.61 (t, *J* = 7.0 Hz, 5H), 3.53 (t, *J* = 5.1 Hz, 2H), 3.36 – 3.24 (m, 2H), 2.85 (m, 4H), 2.46 (t, *J* = 6.4 Hz, 3H), 1.44 (s, 18H).

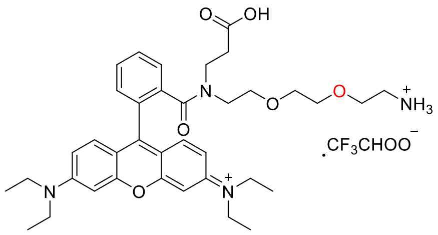

To a solution of Rhodamine B (1 g, 2.09 mmol) in DMF (10 mL) was added triethylamine (0.87 mL, 6.27 mmol) and HBTU (0.87g, 2.30 mmol). After 10 minutes, **S2** (0.87 g, 2.30 mmol was added and the reaction mixture was stirred at room temperature for 2 h. After complete reaction, 50 mL ethyl acetate was added into the reaction flask, followed by washing with water (50 ml, 2 x) and brine (50 mL). The organic layer was dried over MgSO4 and evaporated to get the intermediate **S3**, of sufficient purity for further use.

To a solution of the intermediate **S3** in DCM (2 mL) was added TFA (2 mL) and the reaction mixture was stirred overnight at room temperature. After the reaction finished, evaporated all solvents to get the product (red solid, 60%). $^1$H NMR (600 MHz, MeOD-$d_4$) δ 7.63 – 7.46 (m, 3H), 7.34 – 7.27 (m, 1H), 7.10 (dd, *J* = 9.5, 6.5 Hz, 2H), 6.89 – 6.82 (m, 2H), 6.79 (d, *J* = 2.4 Hz, 1H), 6.75 (d, *J* = 2.3 Hz, 1H), 3.51 – 3.43 (m, 11H), 3.38 (dd, *J* = 9.5, 4.9 Hz, 2H), 3.28 (m, 2H), 3.09 (dt, *J* = 3.2, 1.6 Hz, 9H), 1.10 (t, *J* = 6.4 Hz, 12H).

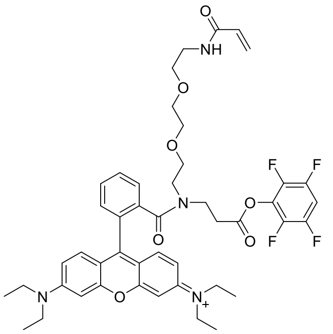

To a solution of the intermediate (0.757 g, 1.11 mmol) in MeOH: THF (v:v = 2 mL: 3 mL) was added a solution of $Na_2CO_3$ (0.239 g, 2.25mmol) in water (3.5 mL) under the protection of $N_2$, followed by cooling the reaction flask to 0°C with an ice bath. A solution of acryloyl chloride (90 µL, 1.11 mmol) in dry dioxane (0.55 mL) was added dropwise to the reaction flask, and the reaction mixture was allowed to come to room temperature over 20 min. After complete reaction, all solvents were evaporated and the residue was used to perform the next step without further purification. To a solution of the intermediate (0.50 g, 0.68 mmol) in ACN (3 mL) was added DCC (0.14 g,0.71 mmol) and 2,3,5,6-Tetrafluorophenol (0.118 g, 0.71 mmol), successively. The reaction was stirred at room temperature under the protection of $N_2$ for 2 h. After the reaction finished, the solution was filtered to remove filtration. Then, the solvent was removed under reduced pressure and the residue was purified by column chromatography to yield the product **S5** as a red solid. $^1$H NMR (600 MHz, DMSO-$d_6$) δ 8.28 – 8.10 (m, 1H), 7.82 – 7.64 (m, 3H), 7.53 (dd, *J* = 5.7, 3.1 Hz, 1H), 7.13 (dd, *J* = 17.5, 10.0 Hz, 4H), 6.98 – 6.94 (m, 2H), 6.87 (s, 1H), 6.24 (ddd, *J* = 15.1, 10.2, 4.8 Hz, 1H), 6.07 (d, *J* = 17.1 Hz, 1H), 5.57 (dd, *J* = 10.2, 2.0 Hz, 1H), 3.72 – 3.60 (m, 8H), 3.51 (m, 2H), 3.47 (t, *J* = 5.9 Hz, 2H), 3.31 (m, 6H), 3.25 (m, 4H), 1.21 (t, *J* = 7.0 Hz, 12H).

General procedure 2: Preparation of small-molecule multifunctional linkers: Pacific Blue-Phalloidin Linker

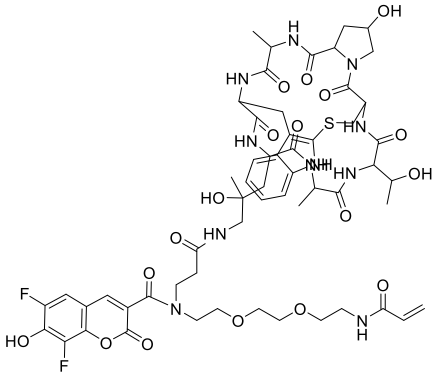

To a solution of the active ester intermediate Pacific blue conjugated with TFP and acryloyl amide (0.1 µmol) (General procedure 1) in DMSO (20 microliter) was added phalloidin amine (tosylate) one equivalent, as 2 mM solution in DMF), and the resulting reaction mixture was stirred at 30$^o$ C for 2h. Upon indication of complete reaction, the crude mixture was purified using preparative HPLC, and the desired products **S6** was used as such.

General procedure 3: Preparation of small-molecule multifunctional linkers: Maleimido linker

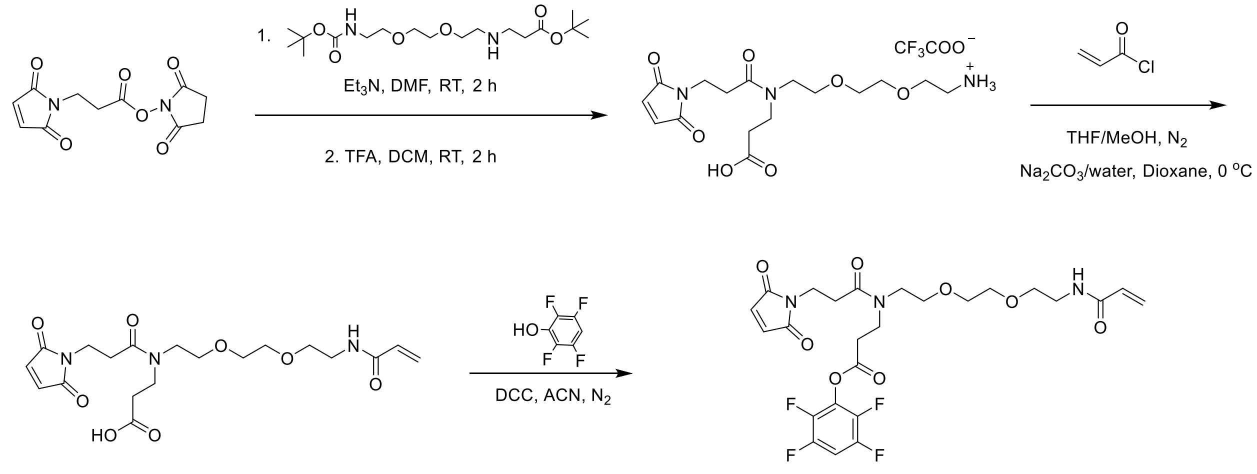


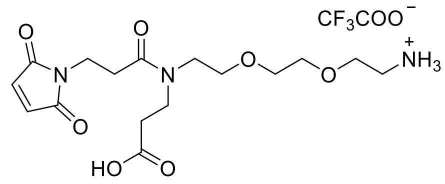

To a solution of Succinimidyl 3-maleimidopropionate (0.32 g, 1.20 mmol) in DMF (4 mL) was added triethylamine (0.35 mL, 2.50 mmol) and **S2**. The reaction mixture was stirred at room temperature for 2 h. After complete reaction, 50 mL ethyl acetate was added into the reaction flask, followed by washing with water (50 mL, 2 x) and brine (50 mL). The organic layer was dried over $MgSO_4$ and evaporated to get the intermediate without further purification. To a solution of the intermediate in DCM (2 mL) was added TFA (2 mL) and the reaction mixture was stirred overnight at room temperature. After the reaction finished, evaporated all solvents to obtain the product **S7** as a (light yellow oil, 90%). $^{1}$H NMR (300 MHz, MeOD-$d_4$) δ 6.83 (s, 2H), 3.65 (m, 14H), 3.17 (m, 2H), 2.80 (m, 2H), 2.64 (m, 2H).

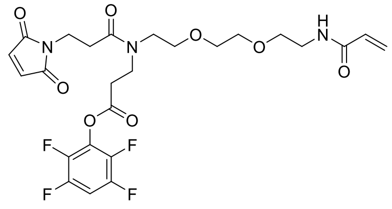

To a solution of the intermediate **S7** (0.458 g, 0.94 mmol) in MeOH: THF (v/v 2:3, 5 mL) was added a solution of $Na_2CO_3$ (0.200 g, 1.89 mmol) in water (3.5 mL) under the protection of $N_2$, followed by cooling the reaction flask to 0°C with an ice bath. A solution of acryloyl chloride (99 µL, 1.23 mmol) in dry dioxane (0.85 mL) was added dropwise to the reaction flask, and the reaction mixture was allowed to come to room temperature over 20 min. After complete reaction, all solvents were evaporated and the residue was used to perform the next step without further purification. To a solution of the crude reaction intermediate (0.40 g, 0.95 mmol) in ACN (5 mL)

was added DCC (0.233 g, 1.14 mmol) and 2,3,5,6-Tetrafluorophenol (0.189 g, 1.14 mmol), successively. The reaction was stirred at room temperature under the protection of $N_2$ for 2 h. After the reaction finished, the solution was filtered to remove filtration. Then, the solvent was removed under reduced pressure and the residue was purified by column chromatography to obtain the desired product **S8** as a clear white oil. $^1$H NMR (300 MHz, MeOD-$d_4$) δ 7.42 (m, 1H), 6.81 (s, 2H), 6.26 (m, 2H), 5.64 (dd, *J* = 9.3, 2.6 Hz, 1H), 3.71 – 3.54 (m, 14H), 3.47 – 3.41 (m, 2H), 3.06 (t, *J* = 6.9 Hz, 2H), 2.81 (dd, *J* = 14.4, 7.1 Hz, 2H).

General procedure 4: Preparation of small-molecule multifunctional linkers: Tetrazine Linker

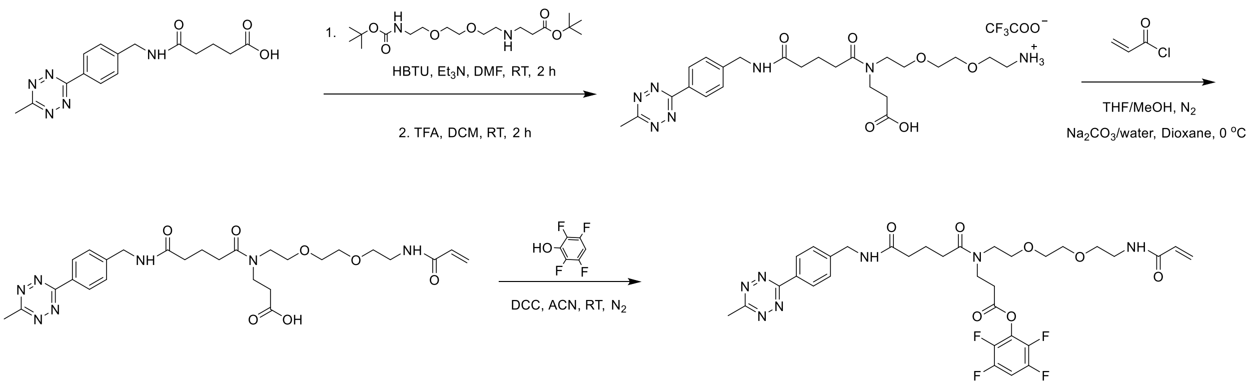


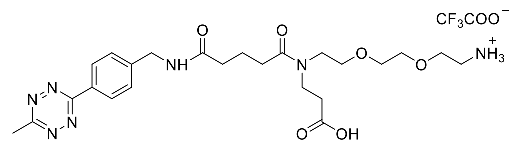

To a solution of the starting material (0.32 g, 1.02 mmol) in DMF (4 mL) was added triethylamine (0.42 mL, 3.04 mmol), HBTU (0.462 g, 1.22 mmol), and **S2**. The reaction mixture was stirred at room temperature for 2 h. After complete reaction, 50 mL ethyl acetate was added into the reaction flask, followed by washing with water (50 mL, 2 x) and brine (50 mL). The organic layer was dried over $MgSO_4$ and evaporated to get the intermediate without further purification. To a solution of the intermediate in DCM (2 mL) was added TFA (2 mL) and the reaction mixture was stirred overnight at room temperature. After the reaction finished, evaporated all solvents to obtain the desired product **S9** as a pink solid (95%).

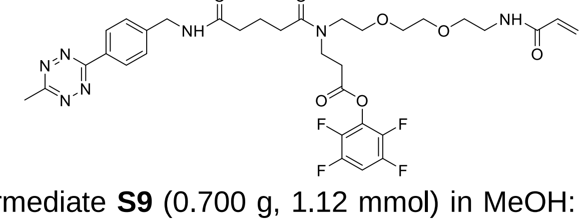

To a solution of the intermediate **S9** (0.700 g, 1.12 mmol) in MeOH: THF (v:v 4:6, 10 mL) was added a solution of $Na_2CO_3$ (0.237 g, 2.24 mmol) in water (3.5 mL) under the protection of $N_2$, followed by cooling the reaction flask to 0°C with an ice bath. A solution of acryloyl chloride (90

μL, 1.12 mmol) in dry dioxane (0.75 mL) was added dropwise to the reaction flask, and the reaction mixture was allowed to come to room temperature over 20 min. After complete reaction, all solvents were evaporated and the residue was used to perform the next step without further purification. To a solution of the intermediate (0.275 g, 0.48 mmol) in ACN (5 mL) was added DCC(0.110 g, 0.53 mmol) and 2,3,5,6-Tetrafluorophenol (0.084 g, 0.51mmol), successively. The reaction was stirred at room temperature under the protection of $N_2$ for 2 h. After the reaction finished, the solution was filtered to remove filtration. Then, the solvent was removed under reduced pressure and the residue was purified by column chromatography to yield the product **S10** as a pink oil. $^1$H NMR (300 MHz, MeOD) δ 7.47 (m, 4H), 6.91 – 6.43 (m, 1H), 6.23 (m, 2H), 5.71 – 5.56 (m, 1H), 4.44 (m,1H), 3.71 – 3.50 (m, 15H), 3.44 (m, 3H), 3.09 (m, 1H), 2.55 – 2.46 (m, 2H), 2.35 (m, 2H).

General procedure 5: Preparation of small-molecule multifunctional linkers: TCO Linker

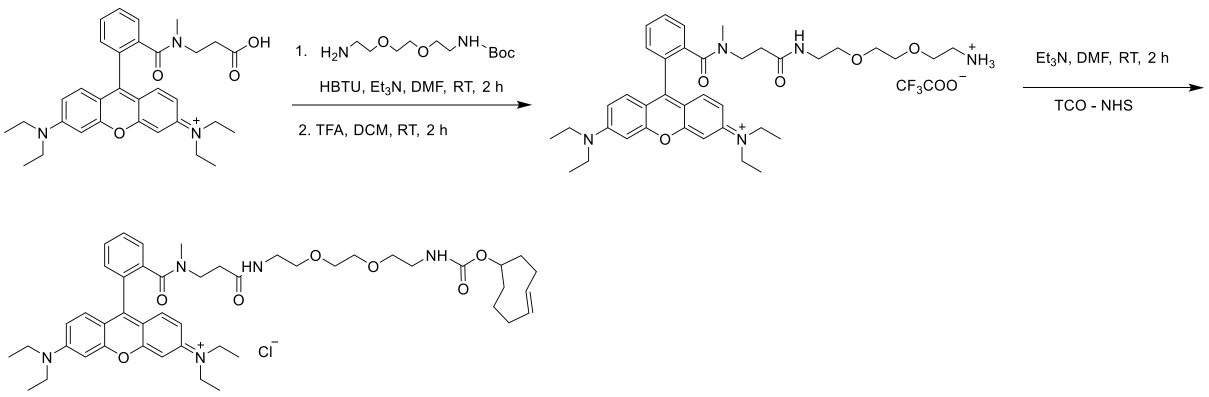


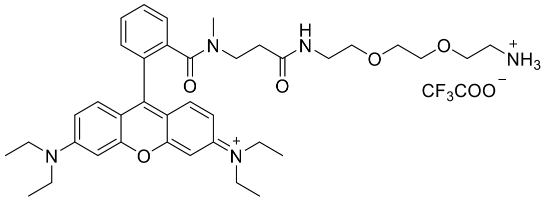


To a solution of the starting Rhodamine B derivative (0.83 g, 1.45 mmol) in DMF (10 mL) was added triethylamine (0.61 mL, 4.39 mmol), HBTU (0.609 g, 1.61 mmol), and N-Boc- 2,2'-(ethylenedioxy)diethylamine (0.407 g, 1.75 mmol), successively. The reaction mixture was stirred at room temperature for 2 h. After complete reaction, 50 mL ethyl acetate was added into the reaction flask, followed by washing with water (50 ml, 2 x) and brine(50 mL). The organic layer was dried over $MgSO_4$ and evaporated to get the intermediate without further purification. To a solution of the intermediate in DCM (2 mL) was added TFA (2 mL) and the reaction mixture was stirred overnight at room temperature. After the reaction finished all solvents were evaporated to yield the product **S11** in sufficient purity for further reaction (Purple solid, 95%).

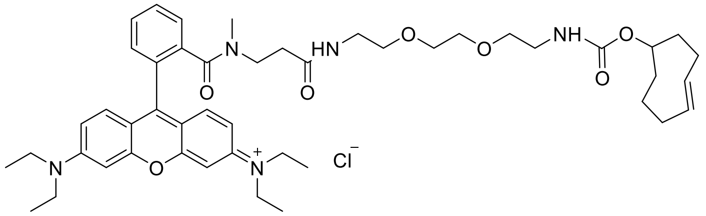


To a solution of the intermediate (37 mg, 56 µmol) in DMF (2 mL) was added triethylamine (19.5 µL, 140 µmol) and TCO-NHS (15 mg, 56 µmol). The reaction mixture was stirred at room temperature for 1 h. After complete reaction, the solvent was evaporated and the residue was purified by column chromatography to get the target compound **S12** (red solid, 56%).$^{1}$H NMR (600 MHz, MeOD-$d_4$) δ 7.80 – 7.74 (m, 2H), 7.74 – 7.66 (m, 1H), 7.51 (dd, *J* = 8.4, 6.5 Hz, 1H), 7.30 (t, *J* = 9.4 Hz, 2H), 7.09 (dt, *J* = 9.4, 2.8 Hz, 2H), 6.98 (d, *J* = 2.4 Hz, 2H), 5.69 (dd, *J* = 16.9, 9.0 Hz, 1H), 5.66 – 5.59 (m, 1H), 4.73 – 4.60 (m, 1H), 3.75 – 3.69 (m, 8H), 3.59 (dd, *J* = 11.5, 6.4 Hz, 3H), 3.51 (dd, *J* = 9.6, 3.9 Hz, 2H), 3.43 (t, *J* = 7.0 Hz, 2H), 3.23 (q, *J* = 7.3 Hz, 12H), 2.11 (t, *J* = 7.0 Hz, 2H), 1.93 – 1.80 (m, 2H), 1.63 (dd, *J* = 13.1, 9.1 Hz, 1H), 1.60 – 1.54 (m, 1H), 1.33 (d, *J* = 7.3 Hz, 24H, including residual solvent peaks).

General procedure 6: Preparation of small-molecule multifunctional linkers: Pacific Blue-lipid Linker

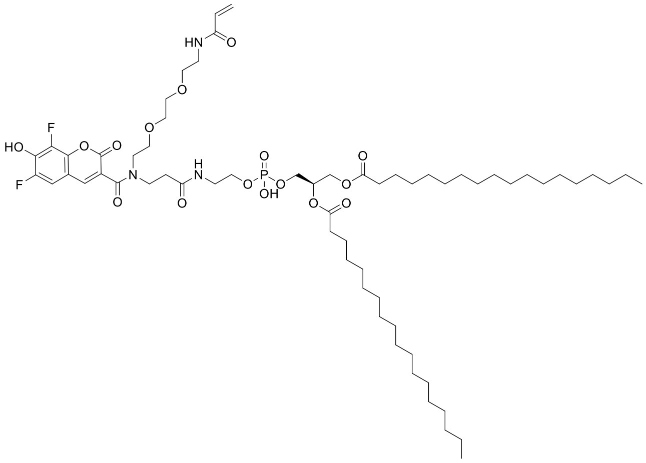


To a solution of the active ester intermediate Pacific blue conjugated with TFP and acryloyl amide (14 mg, 0.022 mmol) (General procedure 1) in Chloroform: water (v:v, 5:1, 6 mL) was added triethylamine (6 µL, 0.043 mmol) and 1,2-Distearoyl-sn-glycero-3-phosphoethanolamine (17.8 mg, 0.024 mmol), and then the resulting reaction mixture was stirred overnight at room temperature. Upon indication of complete reaction, the crude mixture was purified by column chromatography, and the desired products **S13** was obtained as a pale yellow film. Other analogues were prepared following this general procedure. $^{1}$H NMR (300 MHz, MeOD-$d_4$) δ 7.87 (s, 1H), 7.02 (d, *J* = 10.6 Hz, 1H), 6.39 – 6.10 (m, 2H), 5.63 (dd, *J* = 9.6, 2.2 Hz, 1H), 5.22 (brs, 1H), 4.45 (d, *J* = 11.8 Hz, 1H), 4.19 (dd, *J* = 12.0, 6.6 Hz, 1H), 4.05 – 3.88 (m, 4H), 3.62 (m, 10H), 3.45 (m, 2H), 2.32 (dd, *J* = 12.9, 7.1 Hz, 4H), 1.61 (brs, 4H), 1.47 – 1.11 (m, 64H), 0.90 (t, *J* = 6.4 Hz, 6H).

### Dye Portfolio

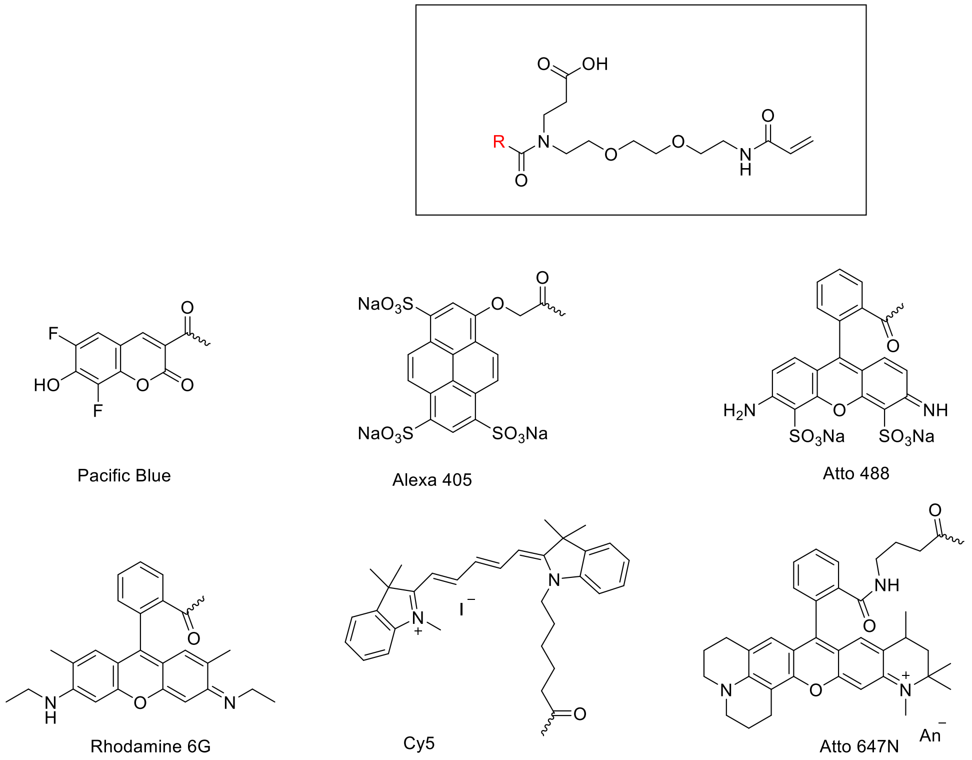


Figure S1. Direct triple linkers with respective dye examples

### Cell culture

HeLa cells were obtained from ATCC and cultured at 37 °C in a 5 % $CO_2$ humidified atmosphere in high glucose (4.5 g/L), glutamine free, phenol red-free Dulbecco's Modified Eagle Medium (DMEM; Life Technologies) supplemented with 10% (v/v) fetal bovine serum, 50 µg/ml gentamicin (Life technologies) and 1% glutamax. When cells reached 70 % confluency, they were washed with 1x DPBS (no calcium, no magnesium; Life Technologies) before detaching with 10x TrypLETM Enzyme (Life Technologies). Afterwards, cells were seeded into the correct imaging chambers depending on the experiment.

### Cytostatics

HeLa cells were seeded into 8-well chambers (Thermo Scientific, 155411) at a density of 25 x $10^3$ cells/well. The next day, cells were fixed in 4% paraformaldehyde (PFA) for 10 minutes and washed 3x for 5 minutes with 1x PBS before permeabilization with 0.2% Triton X-100 at room temperature for 15 minutes. Cells were then washed with 1xPBS followed by at least 1h incubation with 0.25 µM of a multivalent linker with phalloidin-Rhodamine B targetting F-actin. After staining, all cells were washed for 5 minutes with 1x PBS before imaging.

### Immunostaining

HeLa cells were seeded onto size 22 mm x 22mm #1.5 coverglasses at a density of 40 x $10^3$ cells/$cm^2$ and cultured as mentioned in the previous section. The next morning, cells were fixed in 4% PFA for 10 minutes and PFA was quenched in $NH_4Cl$ for 10 minutes. Cells were permeabilized with 0.2% Triton X-100 for 15 minutes at room temperature, washed 3x for 5 minutes with 1x PBS and blocked with blocking buffer (10% fetal bovine and 0.1% Tween-20 in 1x PBS) for 15 minutes. Next, coverslips were incubated with primary antibodies (mouse anti α-tubulin, Abcam, ab7291) in blocking buffer at a concentration of 2 ug/mL for 1 hour at room temperature and washed with PBS three times for 5 minutes each. Specimens were incubated with dye-conjugated secondary antibodies (Goat Anti-Mouse IgG, Abcam ab6708) or DNA-conjugated antibodies (Donkey Anti-Mouse IgG, Abcam ab6707) for 1 hour in blocking buffer at a concentration of 10 µg/mL or with a dilution of 1:100 respectively, and washed with blocking buffer three times for 5 minutes each.

### Lipid staining

HeLa cells were cultured into 8-well chambers (Thermo Scientific, 155411) at a density of 25 x $10^3$ cells/well. After 24 hours, cells were fixed in 4% PFA for 10 minutes and washed with 1xPBS before permeabilization with 0.5% Tween-20 at room temperature for 10 minutes. After washing the cells 3 times for 5 minutes with 1x PBS, membranes were stained for 1 h with 50µM trivalent lipids followed by washing for 5 minutes with 1x PBS before imaging.

### RNA FISH

#### Probe preparation

All DNA oligos were dissolved in deionized water and RNA oligos in RNAse free water at concentrations between 0.1–1 mM and stored at -20 °C. To link the amine modified tiling probes to the amine-modified initiators for ROLLFISH and smHCR experiments, equal molar amounts of each of the tiling probes were reacted with 5 times molar amount of TCO-NHS. Amine-modified initiators were reacted with 5 times molar amount of the trivalent linker **S10**. The reactions were carried out overnight at 4 °C in 1x PBS in a total volume of 1.2 mL. After the reaction, resulting functionalized probes were purified using ethanol precipitation. In brief, 0.5 volumes of 3 M sodium acetate and 2.5 volumes of 100% ethanol were added and the mixture was kept at -80 °C for 2 hours. Next, the mixture was spun down at 4 °C, 16000 g for 30 min. Supernatant was discarded and the clearly visible pellet was washed with 100% ethanol twice, each time spinning down at 4 °C, 16000 g for 2 min. The pellets were then air dried and resuspended in 1x PBS at a concentration of 1 mM. To form the final RNA FISH probes, tiling probes were mixed at equimolar amounts with the initiators for 1 hour at RT. No further purification steps were performed.

#### RollFISH

Cells were rinsed in 1x PBS and then fixed in 4% PFA in PBS for 20 minutes and washed two times in 1x PBS for 5 minutes. Samples were then dried in increasing order in 70%, 85% and 100% ethanol for 5 minutes each. Samples were stored at -80 °C until further use. After incubating for 10 minutes in wash buffer (30% formamide, 5x SSC, 0.1% Tween-20), the samples were incubated overnight at RT in freshly prepared tiling buffer consisting of 30% formamide, 5x SSC,

100 µg/mL salmon sperm DNA, 0.25 mg/mL tRNA, 2 mM RVC and 0.1 µM of each of the 24 tiling probes. The next day the samples were washed three times for 10 minutes in wash buffer. Then the samples were incubated in freshly made padlock buffer (20% formamide, 20 mM TrisHCl, 75 mM KCl, 10 mM MgCl2, 0.5 mM NAD, 0.01% Triton X-100, 100 nM padlock probe, 50 µM each dNTP, 0.5 enzyme units/µL Ampligase; pH 8.3) for 30 minutes at 37 °C followed by 45 minutes' incubation at 45 °C. Then samples were washed twice times in PBST (1x PBS, 0.1% TWEEN-20 pH 7.4) for 5 minutes followed by incubation in freshly made rolloni formation buffer (1 enzyme unit/µL phi29 polymerase, 1x phi29 polymerase reaction buffer, 0.25 mM of each dNTP, 0.2 µg/µL BSA, 5% glycerol) for 1 h. The samples were then washed again in PBST two times for 5 minutes. Samples were subjected to a final incubation in freshly made labeling buffer (2X SSC, 20% formamide, 100 nM detection probe) for 30 minutes at 37 °C. Finally, the samples were washed three times in PBST for 5 min. Samples were kept at 4 °C in 1x PBS until imaging.

#### smHCR

Cells were rinsed in 1x PBS and then fixed in 4% PFA in 1x PBS for 15 minutes and washed three times in 2x SSC for 5 minutes. Samples were stored in 70% ethanol at -20 °C until further use. Samples were dried, then rehydrated in 2x SSC for 5 minutes and then subjected to overnight incubation at 37 °C in freshly mixed hybridization buffer consisting of 10% formamide, 10% dextran sulfate, 2x SSC and 1 to 24 nM of the tiling probe mix containing 24 oligo tiles at equimolar concentrations. On the next day samples were washed for 15 minutes in wash solution (30% formamide, 2x SSC, 0.1% TritonX-100), then rinsed 5 times with 2x SSC. Then samples were incubated for 45 minutes in freshly mixed amplification buffer (10% dextran sulfate, 2x SSC, 120 nM of both h1 and h2 B3-HCR-amplifier), rinsed again 5 times with 5x SSC and kept in freshly made enzymatic anti bleaching buffer (20 mM TrisHCl, 50 mM NaCl, 40 mM glucose, 12 mM trolox, pyranose oxidase (3 enzyme units/mL), catalase (90 enzyme units/mL), pH 8) at 4 °C until imaging.

#### Gelation, digestion, expansion and post-expansion labeling

Gelation solution (1 x PBS, 2 M NaCl, 8.625% (w/w) sodium acrylate, 2.5% (w/w) acrylamide and 0.15% (w/w) N,N'-methylenebisacrylamide enriched with 0.15% tetramethylenediamine, 0.15% ammonium persulfate and 0.01% 4-hydroxy-TEMPO) was prepared and kept on ice until further use to prevent premature gelation. A gelation chamber was prepared by placing two size 22 mm x 22 mm #1.5 coverslips on a Sigmacote® (Sigma Aldrich) treated glass slide spaced by ± 1.5 cm. Next, cells were washed with the gelation solution, gelation solution was removed again and a 80 µl droplet of gelation solution was placed in between the two size 22 mm x 22 mm #1.5 coverslips. The coverslip containing the sample was then introduced on top of the gelation chamber, cells facing down, and the construction was sealed using 4 binder clips. Gelation took place at 37 °C under $N_2$ atmosphere for 2 hours. After gelation, the Sigmacote® treated glass slide was carefully removed from the gelation chamber using the sharp edge of a razor blade and using the same razor blades, the desired size of the gel was cut out. Gels were transferred to a 6-well plate, making use of the supporting coverslip, and incubated in 2-3 mL of proteinase K (New England Biolabs) diluted to 8 U/mL in digestion buffer (50 mM Tris (pH 8), 1 mM EDTA, 0.5% TritonX-100, 0.8 M guanidine HCl) for 12 h at RT. Fluorescently stained samples were expanded by incubation in an excess of deionized water 4 times for 10 minutes each. Samples stained with DNA-conjugated antibodies were washed 3 times for 30 minutes each with PBS, incubated in hybridization wash buffer (20% formamide in 2x SSC) for 10 minutes and incubated overnight with 100 nM readout oligo in hybridization buffer (20% formamide, 10% dextran sulfate, 0.1% Tween-20 in 2x SSC) in an airtight container at 37°C. After hybridization, gels were washed 1 time for 10 minutes with hybridization wash buffer supplied with 1 ug/mL DAPI and two times for 10 minutes each with hybridization wash buffer. Finally, gels were expanded with 0.5x PBS.

#### Fluorescence imaging

Imaging was performed on an inverted Leica true confocal scanner SP8 X system (Wetzlar, Germany) equipped with a HCPLAPO CS2 63X water immersion objective (NA 1.2). DAPI stainings were imaged using a 405 nm pulsed diode laser. A supercontinuum white light laser (SuperK EXTREME/FIANIUM, NKT photonics, Birkerød, Denmark) was used for the excitation of all other fluorophores and laser light was filtered by a notch filter when the correct wavelength was available. Prism dispersion and spectral detection were used to separate the correct signal picked up by a Leica Hybrid Detector. The laser power of the supercontinuum white light laser, the gain, the pinhole size (1 airy unit (AU)) and the line averaging were kept constant when samples were compared. Occasionally 0.3 - 12.0 ns gating was applied to minimize reflection when imaging close to the coverslip. Leica Application Suite X was used to acquire the images which were post-processed using Fiji and Huygens Professional (Scientific Volume Imaging b.v.). All images were collected using Nyquist sampling theorem.
It should be noted that, in line with fluorophore dilution in expansion, the strong decrease of fluorescence intensity per volume in expansion can make the use of high laser power (e.g. 100-fold increase 1.84µW to 188µW) unavoidable in certain cases.

#### Antibody-Oligo Conjugates

Antibody-Oligo conjugates were prepared using standard procedures (See Hermanson, Bioconjugate Techniques). For example, encoding oligo ($NH_2$ terminated, 2 mM in PBS (pH 7.2), 50 µL, 100 nmol)) was reacted with **S8** (10 equivs, as a DMSO solution) by incubating at 37°C for 60 minutes. After complete reaction, the DNA was purified through ethanol precipitation (See above). The dried DNA pellet is resuspended in 1x PBS (pH 7.2) and stored at -20°C. Simultaneously, the desired antibody is reacted with 2-iminothiolane (5-20 equivalents, 60 minutes at RT) and after reaction the antibody was purified in 1x PBS using 0.5 ml 40 kDa Zeba desalting columns (ThermoFisher #87766). The activated antibody is combined with the functionalized oligo in the desired ratio (e.g. 1:40) and reacted for 60 minutes at room temperature, followed by purification and concentration using Amicon Ultra 0.5 mL 50 kDa Centrifugal Filters (EMD Millipore #UFC510096). The functionalized antibodies are stored at 4°C. Validation of oligo-antibody conjugation was conducted through denaturing SDS-PAGE gel assay by denaturing antibodies in LDS sample buffer (ThermoFisher #NP0007) with reducing agent (50 mM DTT) at 70°C for 10 minutes. Next, samples were run on a NuPage 4-12% Bis-Tris PAGE gel (ThermoFisher #NP0322PK2) at 200V for 50 minutes. The gels were stained in a solution containing 0.1% (w/v) Coomassie® Brilliant Blue R-250 (VWR chemicals) in 40% (v/v) ethanol and 10% (v/v) acetic acid for 30 minutes and destained in 5% ethanol and 3.5% acetic acid overnight or until the desired background was achieved.

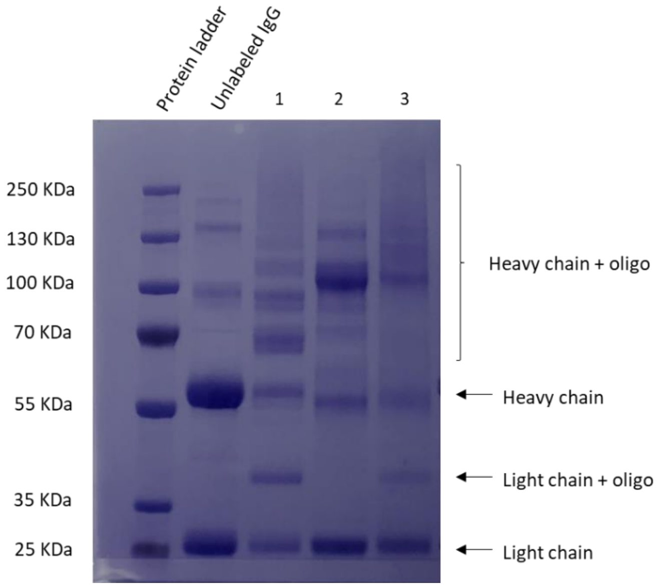


**Supplementary Figure 1 | Visualization of DNA-conjugated antibodies using an SDS-PAGE gel assay.** Lane 1 shows an IgG coupled to barcode sequence 1 without making use of the trivalent linker moiety. Lane 2 and 3 show an IgG (Donkey anti Rabbit and Donkey anti mouse respectively) coupled to encoding probe 1.

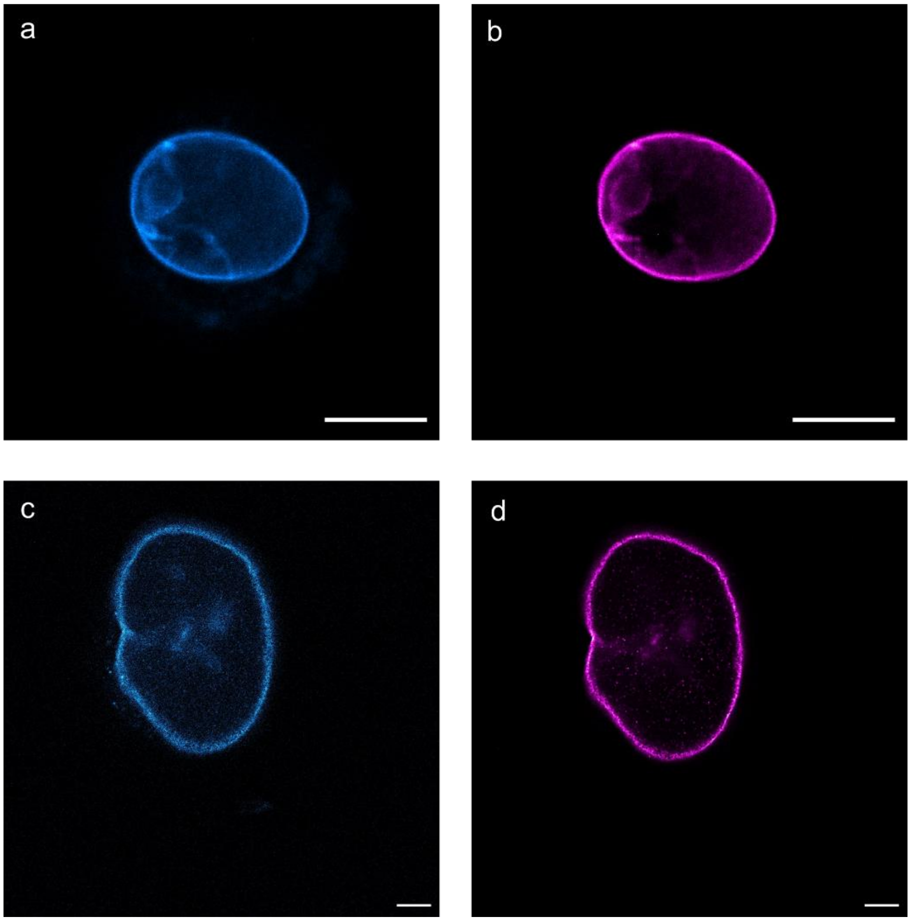


**Supplementary Figure 2 | Validation of primary antibody direct grafting.** Colocalization of Primary antibody anti-Lamin, stained with Pacific Blue, overlapping with fluorescent secondary antibody Atto488. (a) pre-expansion lamin staining in pacific blue channel (b) pre-expansion lamin staining in Atto488 channel (c) post-expansion lamin staining in pacific blue channel (d) post-expansion lamin staining in Atto488 channel (d). Scale bars: 25 µm

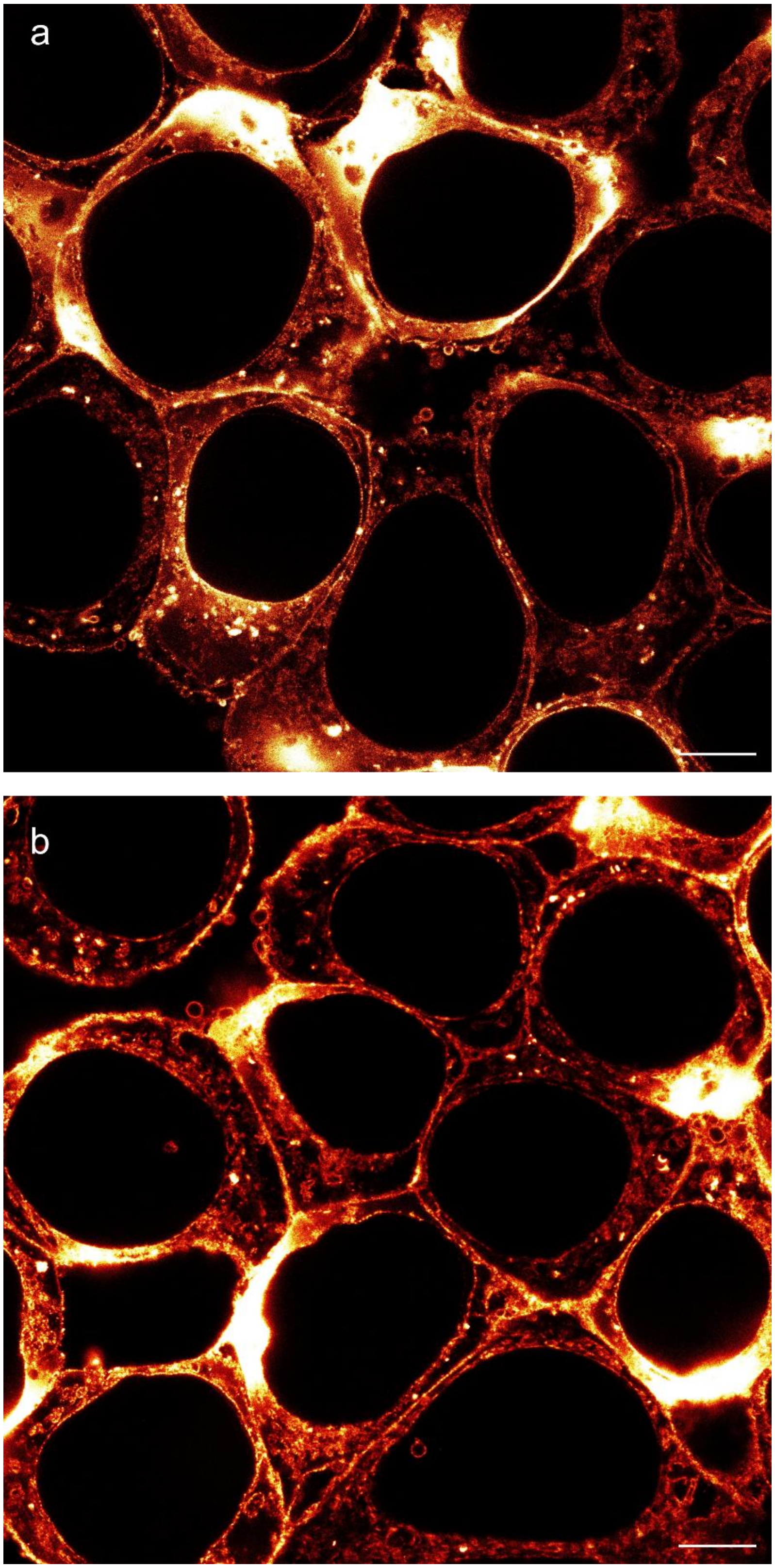


**Supplementary Figure 3 | Post-Expansion images of cellular phospholipid membranes:** a-b) lipid conjugation to fluorescent trivalent linker (Pacific Blue, DSPE, post-expansion) in different regions of the same sample. Scale bars: 25 µm.

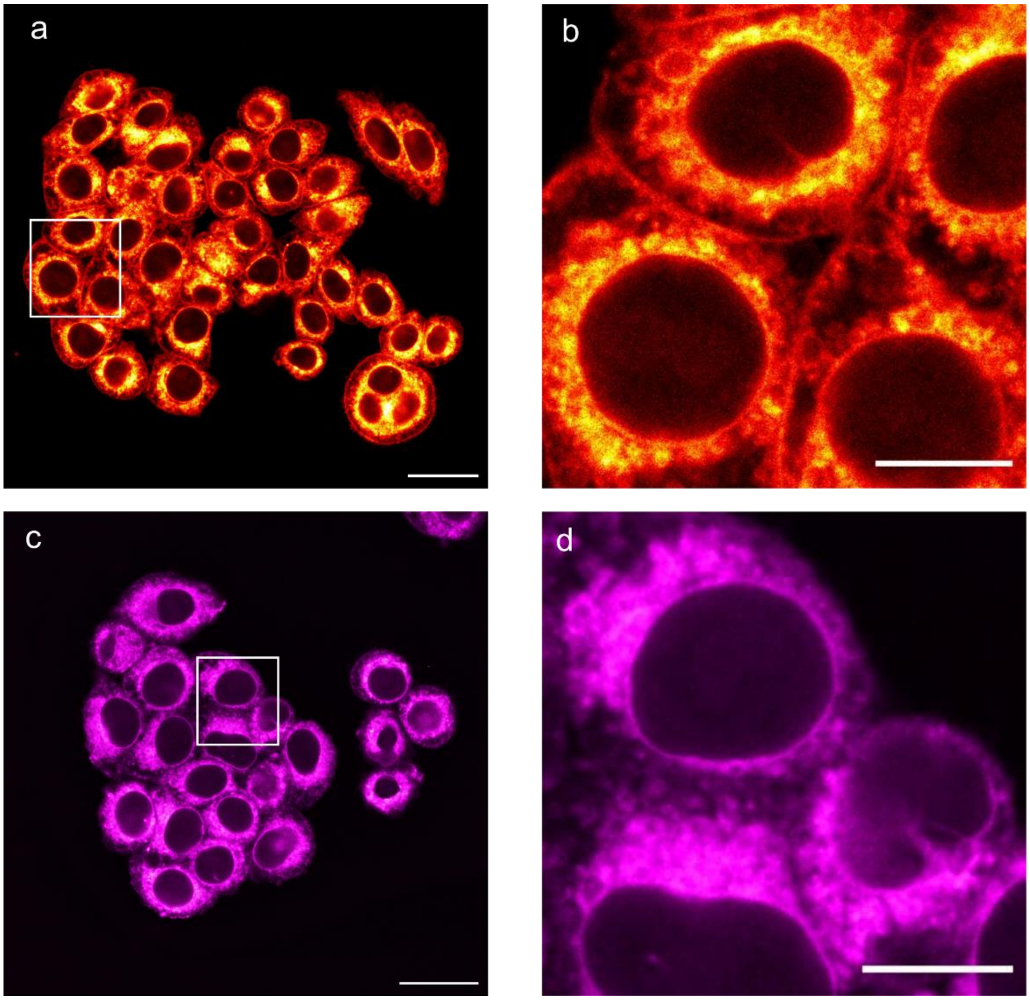


**Supplementary Figure 4 | Lipid conjugates with different dye.** a-b) lipid conjugation to trivalent linker with Atto488 dye (a) pre-expansion image with zoom of the boxed region (b). c-d) lipid conjugation to trivalent linker with Rhodamine 6G dye (c) pre expansion image with zoom of the boxed area (d). Scale bars: 25 µm (a, c), 10µm (b,d).

**Suppelementary Table 1: DNA oligo sequences used.**

| Name | Sequence |
|---|---|
| gusb_tile1 | /5AmMC6/TATATACCATTGCTGCTCGAAACA |
| gusb_tile2 | /5AmMC6/TATAAGCCAATAAAGTCCCGAAGA |
| gusb_tile3 | /5AmMC6/TATATCTCATGTCGGTATCTTGGA |
| gusb_tile4 | /5AmMC6/TATACACTGTTGATCCTCAACACA |
| gusb_tile5 | /5AmMC6/TATACTGGACCAGCTTGCTAATGA |
| gusb_tile6 | /5AmMC6/TATATGCCACCCTCATCCAAAAGA |
| gusb_tile7 | /5AmMC6/TATAACTGGGAACCTGAAGTTGAA |
| gusb_tile8 | /5AmMC6/TATAGCTCATGCATCAGGTAAGGA |

| | |
|---|---|
| gusb_tile9 | /5AmMC6/TATAAAGGAGTACATGTAGGCTGA |
| gusb_tile10 | /5AmMC6/TATAGTGTAGTAGTCAGTCACAGA |
| gusb_tile11 | /5AmMC6/TATACTGTTCGAATCCCGATAGGA |
| gusb_tile12 | /5AmMC6/TATAGAAGGGCTTCCCGTTTATGA |
| gusb_tile13 | /5AmMC6/TATAGTACGAAAGGAATTTGCCCA |
| gusb_tile14 | /5AmMC6/TATACCTCTGAGTAGGGATAGTGA |
| gusb_tile15 | /5AmMC6/TATAATCGGTCACAGAGCTGAAGA |
| gusb_tile16 | /5AmMC6/TATAGAAGTGACTCGTTGCCAAAA |
| gusb_tile17 | /5AmMC6/TATAAACCGCAGGGTGATTTTTGA |
| gusb_tile18 | /5AmMC6/TATAGATAACATCCACGTACGGGA |
| gusb_tile19 | /5AmMC6/TATAAGAACAGCCTTCTGGTACTA |
| gusb_tile20 | /5AmMC6/TATACGAAATTCCAGATGAGCTCA |
| gusb_tile21 | /5AmMC6/TATAAAGTTTTGGGCTGTCTCTGA |
| gusb_tile22 | /5AmMC6/TATATCCGAAACACTGGGTTCTCA |
| gusb_tile23 | /5AmMC6/TATATGAGTCTGCAGTGAGGTAGA |
| gusb_tile24 | /5AmMC6/TATAACACGCACTCCATTTTAGGA |
| smHCR_initiator | /5AmMC6/TATAAAAGTCTAATCCGTCCCTGCCTCTATATCTCCACTC |
| HER2_tile1 | /5AmMC6/TATAATATCCTGCAGGAAGGACAGGCTGGCATTG |
| HER2_tile2 | /5AmMC6/TATATCACTTGGTTGTGAGCGATGAGCACGTAGC |
| HER2_tile3 | /5AmMC6/TATAATAGTTGTCCTCAAAGAGCTGGGTGCCTCG |
| HER2_tile4 | /5AmMC6/TATAGTATTGTTCAGCGGGTCTCCATTGTCTAGC |
| HER2_tile5 | /5AmMC6/TATATCCACAAAATCGTGTCCTGGTAGCAGAGCT |
| HER2_tile6 | /5AmMC6/TATATCAGTGTGAGAGCCAGCTGGTTGTTCTTGT |
| HER2_tile7 | /5AmMC6/TATATCAGGCTCTGACAATCCTCAGAACTCTCTC |
| HER2_tile8 | /5AmMC6/TATATCAAACGTGTCTGTGTTGTAGGTGACCAGG |
| HER2_tile9 | /5AmMC6/TATAAGTCACACAGCTGGCGCCGAATGTATACCG |
| HER2_tile10 | /5AmMC6/TATAAGGATCCCACGTCCGTAGAAAGGTAGTTGT |
| HER2_tile11 | /5AmMC6/TATATTCCATCCTCTGCTGTCACCTCTTGGTTGT |
| HER2_tile12 | /5AmMC6/TATAAAGTGCTCCATGCCCAGACCATAGCACACT |
| HER2_tile13 | /5AmMC6/TATAAACTCCTGGATATTGGCACTGGTAACTGCC |
| HER2_tile14 | /5AmMC6/TATAGCAGAAATGCCAGGCTCCCAAAGATCTTCT |
| HER2_tile15 | /5AmMC6/TATATCCAGAGTCTCAAACACTTGGAGCTGCTCT |
| HER2_tile16 | /5AmMC6/TATAGTCCGGCCATGCTGAGATGTATAGGTAACC |
| HER2_tile17 | /5AmMC6/TATAGTCAGCGAGTAGGCGCCATTGTGCAGAATT |
| HER2_tile18 | /5AmMC6/TATAAAGCAGAGGTGGGTGTTATGGTGGATGAGG |
| HER2_tile19 | /5AmMC6/TATAAAGGAACTGGCTGCAGTTGACACACTGGGT |
| HER2_tile20 | /5AmMC6/TATAAAACAGTGCCTGGCATTCACATACTCCCTG |
| HER2_tile21 | /5AmMC6/TATAAAACAGGTCACTGAGCCATTCTGGGGCTGA |
| HER2_tile22 | /5AmMC6/TATATTATAGTGGGCACAGGCCACACACTGGTCA |
| HER2_tile23 | /5AmMC6/TATAATGTAGGAGAGGTCAGGTTTCACACCGCTG |
| HER2_tile24 | /5AmMC6/TATACTGGCATGCGCCCTCCTCATCTGGAAACTT |
| ROLLFISH_initiator | /5AmMC6/TATATTTTCTAGCGTTAGGGTTAGCGAAGCGTTATAGCAGCGACTACGGT |
| ROLLFISH_padlock | ACGCTTCGCTAACCCTACCTCAATGCTGCTGCTGTACTACTGCGTCTATTTAG |

| | |
|---|---|
| | TGGAGCCCGACCTATCTTCTTTACCGTAGTCGCTGCTATA |
| ROLLFISH_detection | /5Alex546N/TATACCTCAATGCTGCTGCTGTACTAC |
| Encoding Probe 1 | /5AmMC6/AAAAGGTCGATGCCCTAATCCACGTGCTTCCCGC |
| Reporting Probe 1 | GCGGGAAGCACGTGGATTAGGGCATCGACC/3Cy5Sp/ |

**Supplementary Graph 1: Dye survival rate.** Dye survival rate was measured as the ration in emission intensity after polymerization reaction.

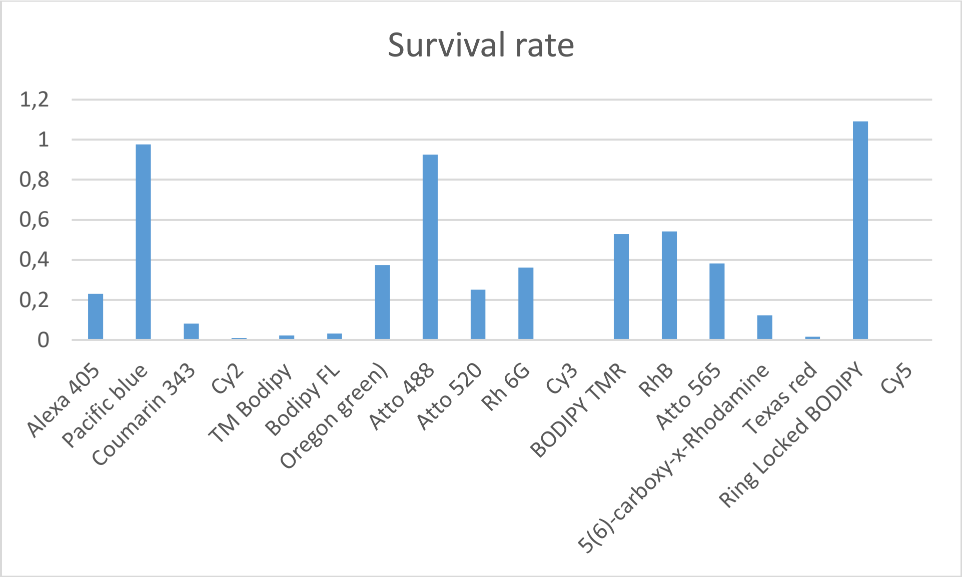
